# pyComBat, a Python tool for batch effects correction in high-throughput molecular data using empirical Bayes methods

**DOI:** 10.1101/2020.03.17.995431

**Authors:** Abdelkader Behdenna, Maximilien Colange, Julien Haziza, Aryo Gema, Guillaume Appé, Chloé-Agathe Azencott, Akpéli Nordor

## Abstract

**Background:** Variability in datasets is not only the product of biological processes: they are also the product of technical biases. ComBat and ComBat-Seq are among the most widely used tools for correcting those technical biases, called batch effects, in, respectively, microarray and RNA-Seq expression data.

**Results:** In this technical note, we present a new Python implementation of ComBat and ComBat-Seq. While the mathematical framework is strictly the same, we show here that our implementations: (*i*) have similar results in terms of batch effects correction; (*ii*) are as fast or faster than the original implementations in R and; (*iii*) offer new tools for the bioinformatics community to participate in its development. pyComBat is implemented in the Python language and is distributed under GPL-3.0 (https://www.gnu.org/licenses/gpl-3.0.en.html) license as a module of the inmoose package. Source code is available at https://github.com/epigenelabs/inmoose and Python package at https://pypi.org/project/inmoose.

**Conclusions:** We present a new Python implementation of state-of-the-art tools ComBat and ComBat-Seq for the correction of batch effects in microarray and RNA-Seq data. This new implementation, based on the same mathematical frameworks as ComBat and ComBat-Seq, offers similar power for batch effect correction, at reduced computational cost.

## 1. Background

Batch effects are the product of technical biases, such as variations in the experimental design or even atmospheric conditions (Lander, 1999; Fare *et al*., 2003). They particularly reveal themselves when merging different datasets, which have likely been built under different conditions. If not corrected, these batch effects may lead to incorrect biological insight, since the variability can be wrongly interpreted as the product of a biological process.

Multiple methods exist that address this problem. They include approaches related to frequentist statistics, such as simple normalization (Yang *et al*., 2002; Tai and Speed, 2012) or principal component analysis (Nielsen *et al*., 2002); and machine learning, such as support-vector machines (Benito *et al*., 2004). One of their main flaws is, however, their incapacity to handle low sample sizes or more than two batches at the same time (Chen *et al*., 2011).

ComBat, originally implemented in the R library sva (Leek *et al*., 2012), is based on the mathematical framework defined in (Johnson *et al*., 2007). This tool leverages a parametric and non-parametric empirical Bayes approach for correcting the batch effect in microarray datasets that works for small sample sizes or in the presence of outliers. Note that the parametric method requires strong assumptions but is largely faster than the non-parametric approach.

ComBat-Seq, also implemented in the R library sva, is based on a similar mathematical framework, where normal distributions are replaced by negative binomial distributions, to better reflect the statistical behavior of RNA-Seq data (Zhang *et al*., 2020).

We introduce in this article pyComBat, a new Python tool implementing ComBat (function pycombat_norm) and ComBat-Seq (function pycombat_seq), following the same mathematical frameworks. Note that the term “pyComBat” implicitly refers to the function pycombat_norm (resp. pycombat_seq) when compared to ComBat (resp. ComBat-Seq). In comparison to both the R implementation and the existing Python implementation of ComBat in the single-cell analysis library Scanpy (Wolf *et al*., 2018), we show that pyComBat yields similar results for adjusting for batch effects in microarray data, but is generally faster, in particular for the usually slow, but more loose, non-parametric method. Similarly, in comparison to the R implementation, we show that pyComBat yields similar results for adjusting for batch effects in RNA-Seq data, but is generally faster. To our knowledge, it is the sole Python implementation of ComBat-Seq.

## 2. Implementation of pyComBat

pyComBat is a Python 3 implementation of ComBat and ComBat-Seq. It mostly uses generic libraries like Pandas (McKinney, 2010) or NumPy (Van Der Walt *et al*., 2011) to mimic ComBat and ComBat-Seq, following the exact same mathematical framework.

Two important features are not directly related to the performance of the software but are of utmost importance. First, pyComBat is available as an open-source software under a GPL-3.0 license, which means anyone can use, modify, distribute and share it. Opening pyComBat to the bioinformatics Python community is the best way for maintaining and improving it, while increasing its robustness. Second, the reliability of pyComBat has been thoroughly checked, using a bench of unit tests (code coverage measured at 88% with Python module “coverage”) serving both as functional tests (to ensure the proper functioning of each submodule) and as non-regression tests (to ease maintenance).

## 3. Results: Comparing pyComBat with ComBat and ComBat-Seq

### a. Datasets used

For software validation, we created two microarray and two RNA-Seq meta-datasets from public data: one on Ovarian Cancer (6 microarray datasets), one on Multiple Myeloma (4 microarray datasets), one on Breast Cancer (originally used for the validation of ComBat-Seq (Zhang *et al*., 2020), 2 RNA-Seq datasets) and one on Colon Cancer (2 RNA-Seq datasets) All meta-datasets are described in more detail in Table 1. We then compared

**Table 1.**
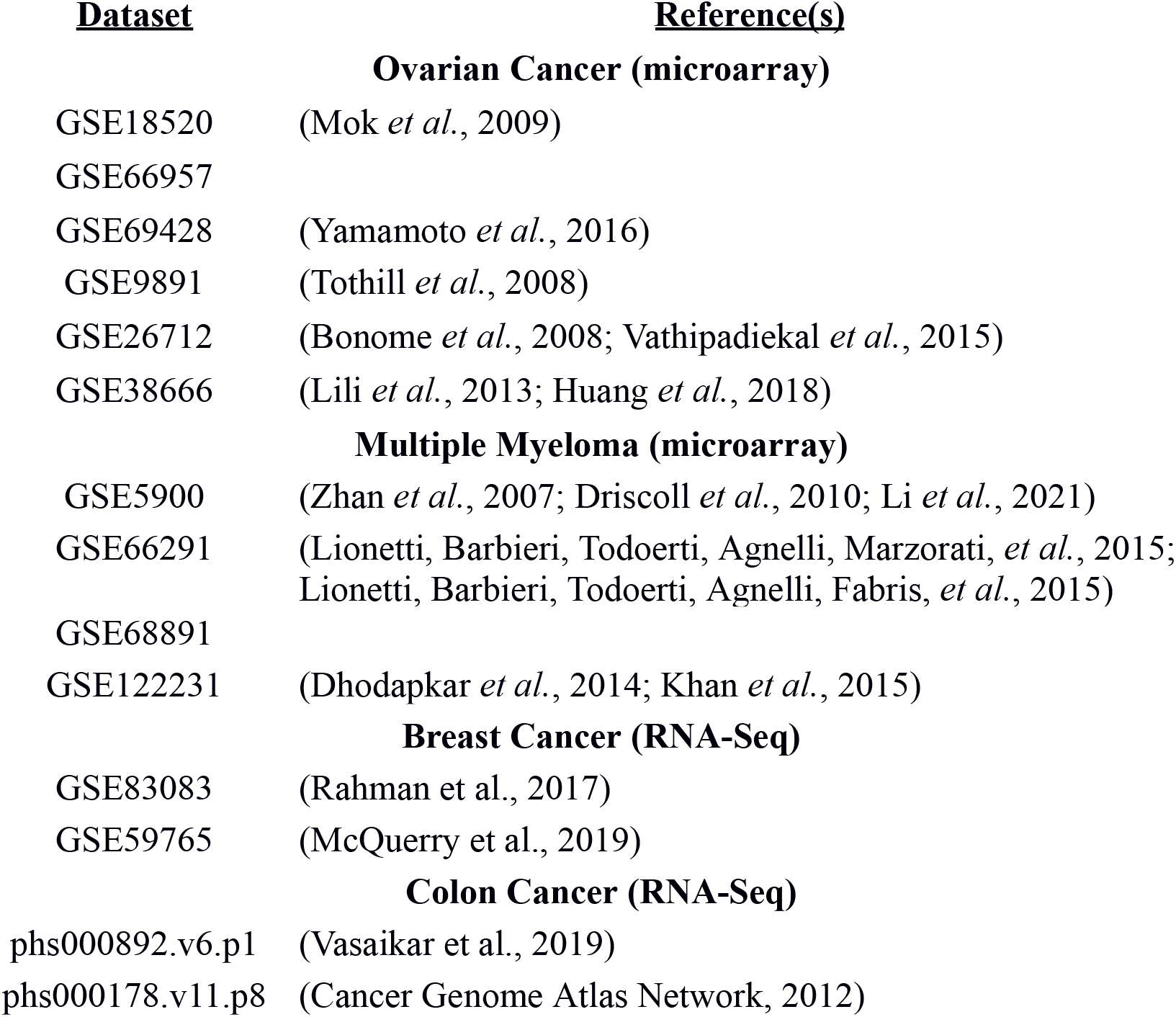
Composition of each meta-dataset used for benchmarking pyComBat, Scanpy’s implementation of ComBat, ComBat and ComBat-Seq.

| <u>Dataset</u> | <u>Reference(s)</u> |
| --- | --- |
| **Ovarian Cancer (microarray)** |  |
| GSE18520 | (Mok *et al.*, 2009) |
| GSE66957 |  |
| GSE69428 | (Yamamoto *et al.*, 2016) |
| GSE9891 | (Tothill *et al.*, 2008) |
| GSE26712 | (Bonome *et al.*, 2008; Vathipadiekal *et al.*, 2015) |
| GSE38666 | (Lili *et al.*, 2013; Huang *et al.*, 2018) |
| **Multiple Myeloma (microarray)** |  |
| GSE5900 | (Zhan *et al.*, 2007; Driscoll *et al.*, 2010; Li *et al.*, 2021) |
| GSE66291 | (Lionetti, Barbieri, Todoerti, Agnelli, Marzorati, *et al.*, 2015; Lionetti, Barbieri, Todoerti, Agnelli, Fabris, *et al.*, 2015) |
| GSE68891 |  |
| GSE122231 | (Dhodapkar *et al.*, 2014; Khan *et al.*, 2015) |
| **Breast Cancer (RNA-Seq)** |  |
| GSE83083 | (Rahman *et al.*, 2017) |
| GSE59765 | (McQuerry *et al.*, 2019) |
| **Colon Cancer (RNA-Seq)** |  |
| phs000892.v6.p1 | (Vasaikar *et al.*, 2019) |
| phs000178.v11.p8 | (Cancer Genome Atlas Network, 2012) |

- ComBat, Scanpy’s implementation of ComBat and pyComBat on the microarray datasets on one hand,
- ComBat-Seq and pyComBat on the RNA-Seq datasets.

for (*i*) efficacy for batch effect correction and (*ii*) computation time.

### b. Batch effect correction

As an implementation of the ComBat and ComBat-Seq algorithms, pyComBat is expected to have similar, if not identical, power in terms of batch effects correction. This is confirmed in Fig.1A, which shows the distribution of differences between the outputs of ComBat and pyComBat, on the Ovarian Cancer dataset (mean = −1.06·10^−7^, 95% CI = [-1.28·10^−3^,1.32 ·10^−4^]), and on Fig. 2A, which shows the distribution of relative differences between the outputs of ComBat-Seq and pyComBat, on the Colon Cancer dataset (mean = 5.4·10^−6^, first non-zero quantile 0.9995). As expected, the differences are distributed closely around zero. The slight variability can be explained by the difference between optimization routines in R and Python (Numpy): while small differences in the fitted distribution parameters have little to no impact for the vast majority of data points, they may be amplified by data adjustment for data points located far away in the distribution tails.

**Fig. 1.**
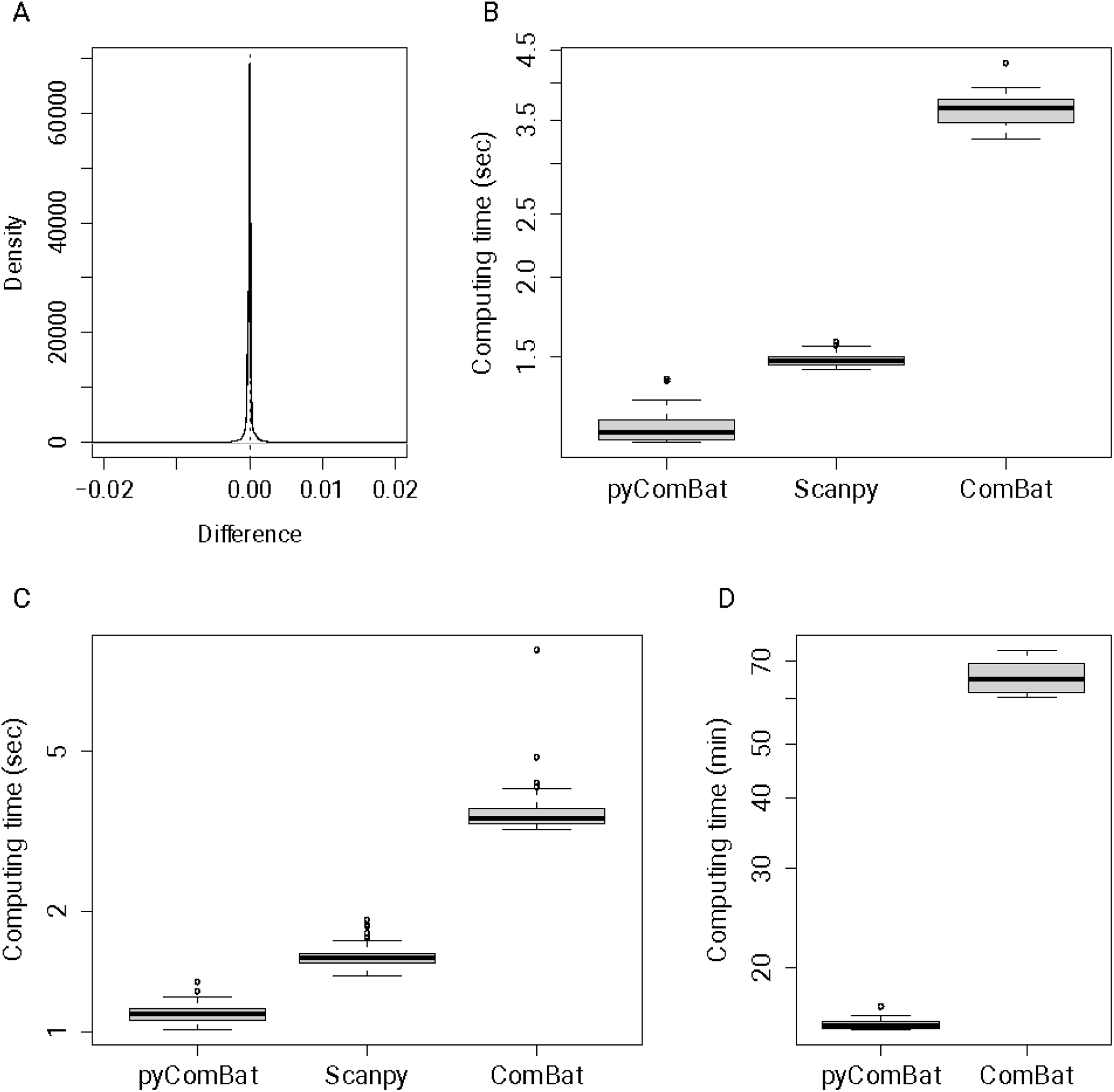
Performance of pyComBat *vs*. Combat *vs*. Scanpy’s implementation of ComBat. **A** Distribution of the differences between the expression matrices corrected for batch effects, respectively by ComBat and pyComBat (parametric version), on the Ovarian Cancer dataset. The vertical dotted line corresponds to zero. **B** Computation time in seconds for pyComBat, Scanpy and ComBat for the parametric method, on the Multiple Myeloma dataset. The y-axis is in a log scale. **C** Computation time in seconds for pyComBat, Scanpy and ComBat for the parametric method, on the Ovarian Cancer dataset. The y-axis is in a log scale. **D** Computation time in minutes for pyComBat (left) and ComBat (right) for the non-parametric method, on the Ovarian Cancer dataset. The y-axis is in a log scale.

**Fig. 2.**
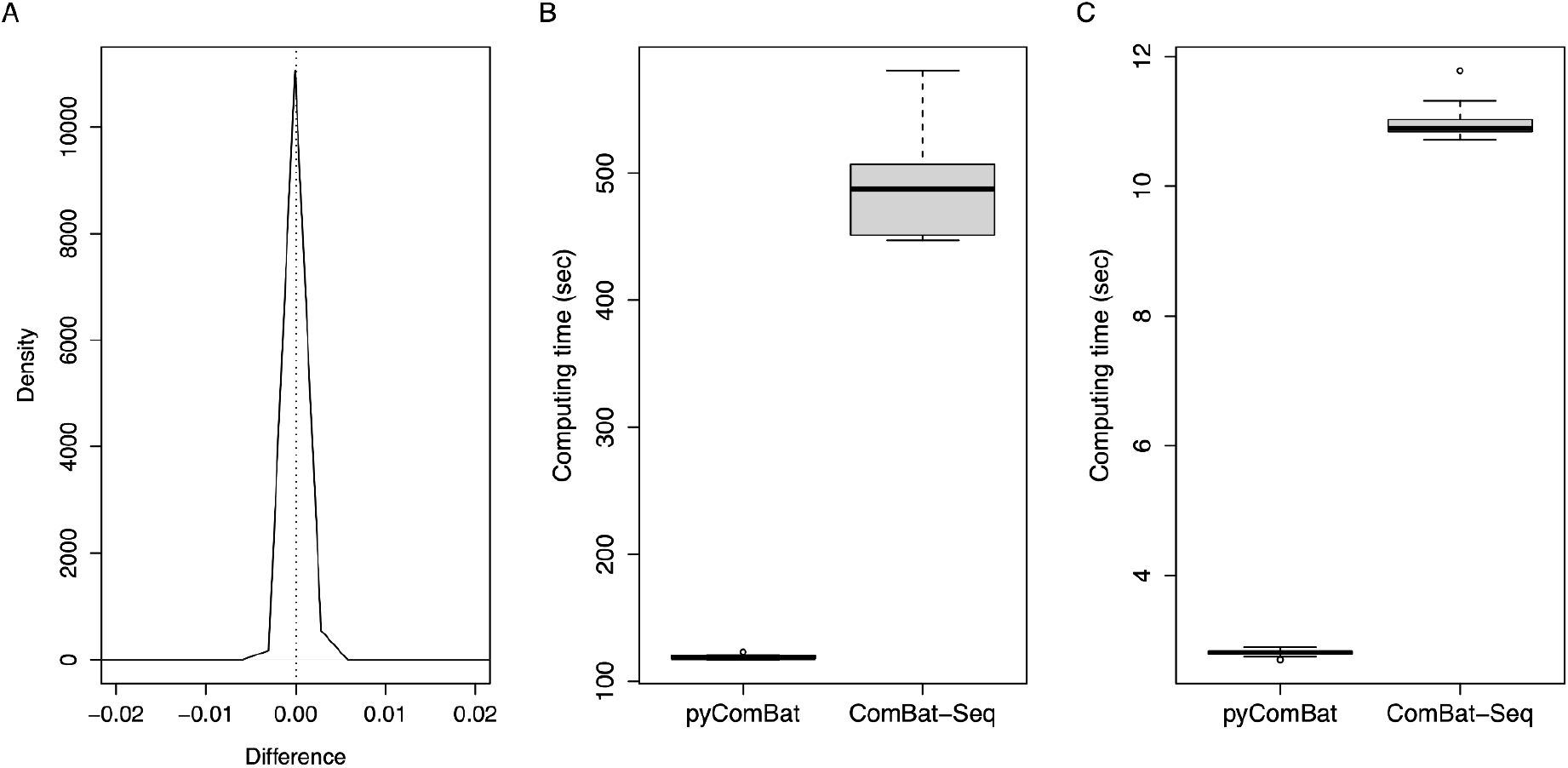
Performance of pyComBat *vs*. ComBat-Seq. **A** Distribution of the relative differences between the expression matrices corrected for batch effects, respectively by ComBat-Seq and pyComBat, on the Colon Cancer dataset. The vertical dotted line corresponds to zero. **B** Computation time in seconds for pyComBat and ComBat-Seq, on the Colon Cancer dataset. **C** Computation time in seconds for pyComBat and ComBat-Seq, on the Breast Cancer dataset.

Additionally, both ComBat-Seq and pyComBat produce the exact same output on the Breast Cancer dataset.

### c. Computation time

Computation time is evaluated by running pyComBat (resp. Scanpy’s implementation of ComBat and ComBat itself) respectively 100 times on both microarray datasets presented in section 3a, with the parametric approach. As Scanpy doesn’t handle the non-parametric approach, only ComBat and pycombat_norm have been tested with it, on the Ovarian Cancer dataset. As for pycombat_seq and ComBat-Seq, they have been run respectively 50 times on the Breast Cancer dataset and 20 times on the Colon Cancer dataset. The reduced number of runs on RNA-Seq datasets is due to the longer, and less variable, computation times.

Owing to Python (Numpy) efficiency in handling matrix operations and matrix manipulations as well as thorough optimization of our code, pyComBat is as fast or even faster than ComBat. The parametric version of the pyComBat performs 4 to 5 times as fast as ComBat, and around 1.5 times as fast as the Scanpy implementation of ComBat, in terms of computation time (fig.1B, fig.1C), on both datasets.

Similar results are observed with the non-parametric version (fig.1D), which is inherently more time consuming, but also less dependent on the distribution of the data. In this case, pyComBat is also approximately 4 to 5 times faster than ComBat, going from more than an hour to around 15 minutes.

Finally, pyComBat appears to be 4 to 5 times faster than ComBat-Seq on both RNA-Seq datasets (Fig. 2B, Fig. 2C).

## 4. Discussion and conclusion

We present pyComBat, a new Python implementation of ComBat and ComBat-Seq, the most commonly used software for batch effects correction on high-throughput molecular data. Our implementation offers the same correcting power, with shorter computation time for the parametric method compared to other implementations, and significantly shorter time for the time-consuming non-parametric version compared to the original R implementations. This reduced computing time opens perspectives for a more generic use of the non-parametric approach to a larger range of datasets.

We have attached importance to making the software open source, coupled with comprehensive documentation. We built a robust set of test cases, in an effort to encourage larger participation from the community. We believe that this will be benefiting the Python bioinformatics community and opening the way towards the translation of other widely used software from R to Python.

## Availability and requirements

**Project name:** InMoose

**Project home page:** https://github.com/epigenelabs/inmoose

**Operating system:** Platform independent

**Programming language:** Python

**Other requirements:** Numpy, Scipy

**License:** GNU GPL3

**Any restrictions to use by non-academics:** None

## Declarations

### Ethics approval and consent to participate

Not Applicable

### Consent for publication

Not Applicable

### Availability of data and materials

The datasets analyzed during the current study are available in the GEO and dbGAP repositories:

https://www.ncbi.nlm.nih.gov/geo/query/acc.cgi?acc=GSE18520

https://www.ncbi.nlm.nih.gov/geo/query/acc.cgi?acc=GSE66957

https://www.ncbi.nlm.nih.gov/geo/query/acc.cgi?acc=GSE69428

https://www.ncbi.nlm.nih.gov/geo/query/acc.cgi?acc=GSE9891

https://www.ncbi.nlm.nih.gov/geo/query/acc.cgi?acc=GSE26712

https://www.ncbi.nlm.nih.gov/geo/query/acc.cgi?acc=GSE38666

https://www.ncbi.nlm.nih.gov/geo/query/acc.cgi?acc=GSE5900

https://www.ncbi.nlm.nih.gov/geo/query/acc.cgi?acc=GSE66291

https://www.ncbi.nlm.nih.gov/geo/query/acc.cgi?acc=GSE68891

https://www.ncbi.nlm.nih.gov/geo/query/acc.cgi?acc=GSE122231

https://www.ncbi.nlm.nih.gov/geo/query/acc.cgi?acc=GSE83083

https://www.ncbi.nlm.nih.gov/geo/query/acc.cgi?acc=GSE59765

https://www.ncbi.nlm.nih.gov/projects/gap/cgi-bin/study.cgi?study_id=phs000892.v6.p1

https://www.ncbi.nlm.nih.gov/projects/gap/cgi-bin/study.cgi?study_id=phs000178.v11.p8

### Authors’ contributions

AB implemented pycombat_norm and compared it with ComBat and Scanpy’s implementation of ComBat. MC implemented pycombat_seq and compared it with ComBat-Seq. AB and MC wrote this manuscript. JH, AG, GA, CAA and AN reviewed code and provided scientific guidance. All authors read and approved the final manuscript.

### Authors’ information

AB and MC should be considered as joint first authors. AG is credited for his contributions while employed at Epigene Labs.

## Acknowledgements

The authors thank Phuong Pham for his advice about the estimation of the efficiency of the adjustments. The authors also thank all the external contributors on the GitHub repository, for their feedback on the code for pyComBat.

## References

Benito, M. et al. (2004) Adjustment of systematic microarray data biases. Bioinformatics, 20, 105–114.

Bonome, T. et al. (2008) A gene signature predicting for survival in suboptimally debulked patients with ovarian cancer. Cancer Res., 68, 5478–5486.

Cancer Genome Atlas Network (2012) Comprehensive molecular characterization of human colon and rectal cancer. Nature, 487(7407), 330–7. doi: 10.1038/nature11252.

Chen, C. et al. (2011) Removing batch effects in analysis of expression microarray data: An evaluation of six batch adjustment methods. PLoS One, 6.

Dhodapkar, M. V. et al. (2014) Clinical, genomic, and imaging predictors of myeloma progression from asymptomatic monoclonal gammopathies (swog s0120). Blood, 123, 78–85.

Driscoll, J.J. et al. (2010) The sumoylation pathway is dysregulated in multiple myeloma and is associated with adverse patient outcome. Blood, 115, 2827–2834.

Fare, T.L. et al. (2003) Effects of atmospheric ozone on microarray data quality. Anal. Chem., 75, 4672–4675.

Huang, C. et al. (2018) Machine learning predicts individual cancer patient responses to therapeutic drugs with high accuracy. Sci. Rep., 8.

Johnson, W.E. et al. (2007) Adjusting batch effects in microarray expression data using empirical Bayes methods. Biostatistics, 8, 118–127.

Khan, R. et al. (2015) Four genes predict high risk of progression from smoldering to symptomatic multiple myeloma (SWOG s0120). Haematologica, 100, 1214–1221.

Lander, E.S. (1999) Array of hope. Nat. Genet., 21, 4.

Leek, J.T. et al. (2012) The SVA package for removing batch effects and other unwanted variation in high-throughput experiments. Bioinformatics, 28, 882–883.

Li, C. et al. (2021) Genetic analysis of multiple myeloma identifies cytogenetic alterations implicated in disease complexity and progression. Cancers (Basel)., 13, 1–15.

Lili, L.N. et al. (2013) Molecular profiling predicts the existence of two functionally distinct classes of ovarian cancer stroma. Biomed Res. Int., 2013.

Lionetti, M., Barbieri, M., Todoerti, K., Agnelli, L., Fabris, S., et al. (2015) A compendium of DIS3 mutations and associated transcriptional signatures in plasma cell dyscrasias. Oncotarget, 6, 26129–26141.

Lionetti, M., Barbieri, M., Todoerti, K., Agnelli, L., Marzorati, S., et al. (2015) Molecular spectrum of BRAF, NRAS and KRAS gene mutations in plasma cell dyscrasias: Implication for MEK-ERK pathway activation. Oncotarget, 6, 24205–24217.

McKinney, W. (2010) Data Structures for Statistical Computing in Python. Proc. 9th Python Sci. Conf., 1697900, 51–56.

McQuerry, J.A., et al. (2019) Pathway activity profiling of growth factor receptor network and stemness pathways differentiates metaplastic breast cancer histological subtypes. BMC cancer, 19.1, 1–14.

Mok, S.C. et al. (2009) A Gene Signature Predictive for Outcome in Advanced Ovarian Cancer Identifies a Survival Factor: Microfibril-Associated Glycoprotein 2. Cancer Cell, 16, 521–532.

Nielsen, T.O. et al. (2002) Molecular characterisation of soft tissue tumours: A gene expression study. Lancet, 359, 1301–1307.

Rahman, M. et al. (2017) Activity of distinct growth factor receptor network components in breast tumors uncovers two biologically relevant subtypes. Genome medicine, 9.1, 1–19.

Tai, Y.C. and Speed, T.P. (2012) A multivariate empirical Bayes statistic for replicated microarray time course data. In, Selected Works of Terry Speed., pp. 617–642.

Tothill, R.W. et al. (2008) Novel molecular subtypes of serous and endometrioid ovarian cancer linked to clinical outcome. Clin. Cancer Res., 14, 5198–5208.

Vathipadiekal, V. et al. (2015) Creation of a human secretome: A novel composite library of human secreted proteins: Validation using ovarian cancer gene expression data and a virtual secretome array. Clin. Cancer Res., 21, 4960–4969.

Van Der Walt, S. et al. (2011) The NumPy array: A structure for efficient numerical computation. Comput. Sci. Eng., 13, 22–30.

Vasaikar, S. et al. (2019) Proteogenomic Analysis of Human Colon Cancer Reveals New Therapeutic Opportunities. Cell. 2019 May 2;177(4):1035–1049.e19. doi: 10.1016/j.cell.2019.03.030.

Wolf, F.A. et al. (2018) SCANPY: Large-scale single-cell gene expression data analysis. Genome Biol., 19, 15.

Yamamoto, Y. et al. (2016) In vitro and in vivo correlates of physiological and neoplastic human Fallopian tube stem cells. J. Pathol., 238, 519–530.

Yang, Y.H. et al. (2002) Normalization for cDNA microarray data: a robust composite method addressing single and multiple slide systematic variation. Nucleic Acids Res., 30, 15e –15.

Zhan, F. et al. (2007) Gene-expression signature of benign monoclonal gammopathy evident in multiple myeloma is linked to good prognosis. Blood, 109, 1692–1700.

Zhang, Y. et al. (2020) ComBat-seq: batch effect adjustment for RNA-seq count data. NAR Genomics Bioinforma., 2.

